# Metacognitive impairments extend perceptual decision making weaknesses in compulsivity

**DOI:** 10.1101/098277

**Authors:** Tobias U. Hauser, Micah Allen, NSPN Consortium, Geraint Rees, Raymond J. Dolan

## Abstract

Awareness of one’s own abilities is of paramount importance in adaptive decision making. Psychotherapeutic theories assume such metacognitive insight is impaired in compulsivity, though this is supported by scant empirical evidence. In this study, we investigate metacognitive abilities in compulsive participants using computational models, where these enable a segregation between metacognitive and perceptual decision making impairments. We examined twenty low-compulsive and twenty high-compulsive participants, recruited from a large population-based sample, and matched for other psychiatric and cognitive dimensions. Hierarchical computational modelling of the participants’ metacognitive abilities on a visual global motion detection paradigm revealed that high-compulsive participants had a reduced metacognitive ability. This impairment was accompanied by a perceptual decision making deficit whereby motion-related evidence was accumulated more slowly in high compulsive participants. Our study shows that the compulsivity spectrum is associated with a reduced ability to monitor one’s own performance, over and above any perceptual decision making difficulty.

## Introduction

Knowing what you did and how well you did it is crucial for achieving one’s goals and making adequate decisions^1^. Humans are burdened with imperfect perception and recollection, and this extends to the metacognitive ability to recognize such deficits. Despite this sub-optimality, we retain an ability to quantify the degree to which we can rely on our behaviour as represented by the feeling of confidence.

Confidence helps us determine how much credit we should assign to an information source, enabling us to calibrate our future behaviour. Metacognitive ability is thus important for good performance, and it is known that metacognitive training improves decision making^2^. However, there are considerable variations in metacognitive performance, i.e. how well humans are able to consciously judge their own performance^3-5^. Poor metacognitive skills, or insight, can have detrimental real-world consequences. For example, one might assign too much credit to a poorly informed decision or exhibit too little trust in a good decision. In extremis, impaired metacognition might lead to systematically bad decisions, for example continuously enacting the same behaviour regardless of outcome, as observed in obsessive checking^6^.

Obsessive-compulsive disorder (OCD) is a condition linked to metacognitive impairment. This disorder is characterized by intrusive thoughts and images (obsessions), and these are coupled to repetitive behaviours (compulsions) which serve to alleviate obsession-induced distress^7^. Initial theories of metacognitive impairments in OCD propose patients overestimate the credibility of their intrusions, believing their likelihood of becoming real^8,9^. Therapy for OCD often targets these (meta-)cognitive biases^6^. More recent accounts propose that metacognitive impairments are not restricted to intrusions, but also apply to memory recollection, although not unequivocally^10–15^. Thus, impairments in meta-memory are believed to drive repetitive checking, because low confidence in one’s own memory is likely to cause a repetition of a previously carried out action. However, findings of lowered confidence in patients with OCD in cognitive domains other than memory^16,17^ suggest OCD patients might suffer from a more general impairment in metacognition.

Traditional studies of metacognition using questionnaires^11,14,18-22^ or subjective confidence ratings^10,12,13,15^ are subject to influences that may mimic a metacognitive impairment, such as systematic response biases in questionnaires and other confidence scales^23^. Here, we define metacognition as the objective sensitivity of confidence ratings to discrimination performance, as defined by signal detection theory. Metacognition thus reflects the degree of insight into one’s behaviour, i.e. how well one knows one’s performance. This model-based measure is robust against general biases in rating behaviours (e.g. generally lower or higher ratings) and is independent of variability in perceptual decision making ability that can directly influence confidence ratings. The latter is of particular importance as OCD patients are reported to suffer from perceptual decision making difficulties^24^. Our model-based computational accounts of metacognition together with controlling task difficulty circumvent these limitations by singling out different contributing factors that influence confidence ratings and metacognition^23^.

In this study, we probed metacognitive abilities along a recently proposed compulsivity spectrum^25,26^ using a perceptual decision making task in two groups of participants with either high or low compulsive scores. These participants were carefully selected from a large cohort so as to match for potential confounding factors, such as depressive or anxiety symptoms^13^. The psychophysical detection task was continuously and automatically adapted for each participant to maintain constant performance levels, allowing us to study separate perceptual decision making and metacognitive differences. Using a hierarchical metacognition model, we analysed participants’ metacognitive efficiency, allowing us to map objective sensitivity of a participant’s subjective beliefs about their performance to actual underlying performance.

Using this computational approach, we found that compulsivity is related to impairments in metacognitive efficiency, difficulties complemented by an independent perceptual decision making impairment.

## Methods

### Participants

We recruited forty participants from a large population-based sample of 2409 young people in London and Cambridge (U-CHANGE study; www.nspn.org.uk^27,28^). We used a directed sampling approach, selecting twenty participants with high compulsivity scores (‘high compulsives’) and twenty participants with low compulsivity scores (‘low compulsives’). For this categorisation we used the PI-WSUR questionnaire^29^ as an index of compulsivity. The groups were selected so as to match in terms of age, gender, depression (using MFQ questionnaire^30^), and anxiety levels (using RCMAS questionnaire^31^). The groups also did not differ in IQ (using vocabulary and matrix subtests of WASI battery^32^) and impulsivity (BIS questionnaire^33^). Participants that reported any psychiatric or neurological disorders were excluded a priori. All participant had normal or corrected-to-normal vision.

The selected groups differed strongly in their compulsivity scores, but were otherwise well matched across all other psychiatric dimensions (Table 1). Two high compulsive participants were excluded from data analysis due to difficulties with the task (staircase failed to converge). The study was approved by the UCL research ethics committee (No. 6218/001) in accordance with the Declaration of Helsinki and all participants gave written informed consent.

**Table 1.**
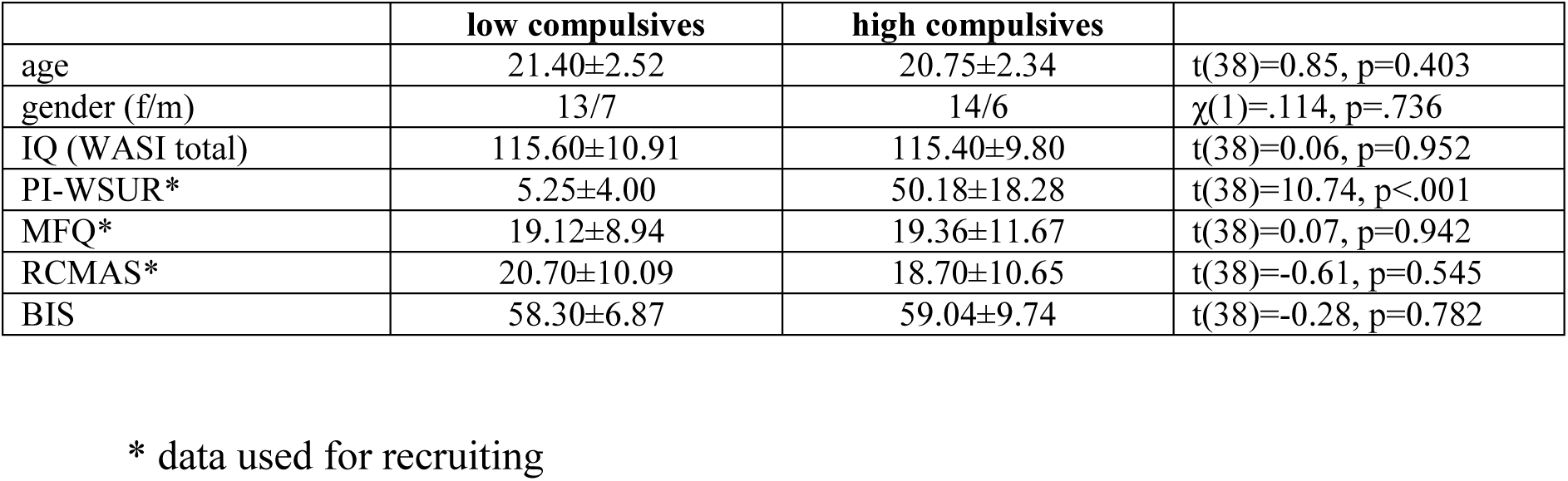
Participants with high and low compulsivity scores. Two groups of participants were recruited from a population-based database, based on their compulsivity scores (PI-WSUR). The groups were matched for other psychiatric dimensions, especially depression (MFQ) and anxiety (RCMAS). Groups did not differ in age, gender, IQ, or impulsivity (BIS). (mean±SD)

|  | **low compulsives** | **high compulsives** |  |
| --- | --- | --- | --- |
| age | 21.40±2.52 | 20.75±2.34 | t(38)=0.85, p=0.403 |
| gender (f/m) | 13/7 | 14/6 | $\chi^2(1)=.114$, p=.736 |
| IQ (WASI total) | 115.60±10.91 | 115.40±9.80 | t(38)=0.06, p=0.952 |
| PI-WSUR* | 5.25±4.00 | 50.18±18.28 | t(38)=10.74, p<.001 |
| MFQ* | 19.12±8.94 | 19.36±11.67 | t(38)=0.07, p=0.942 |
| RCMAS* | 20.70±10.09 | 18.70±10.65 | t(38)=-0.61, p=0.545 |
| BIS | 58.30±6.87 | 59.04±9.74 | t(38)=-0.28, p=0.782 |

\* data used for recruiting

### Task

We used a metacognition task based on a global motion detection paradigm, similar to that reported previously^3,34^. The task (Fig. 1A) consisted of 140 trials subdivided into 10 blocks, with short breaks between blocks. On each trial, participants judged whether the global motion of the randomly moving dots was directed left- or rightwards relative to vertex. Subsequently, participants had to indicate their confidence using a visual analogue scale, where 0 indicated a guess and 100 total certainty. To prevent motor preparation, the starting point of the confidence slider was randomly adjusted to +/-12% of the scale midpoint. Before the main task, participants completed a short training and were also instructed to use the entire confidence scale for their confidence ratings. The task was implemented using Psychtoolbox 3 (www.psychtoolbox.org) in MATLAB (MathWorks Inc.).

The motion signal consisted 1100 black dots presented for 250ms. The motion direction of the dots was determined using a mean motion angle (‘orientation’, in degrees from vertical movement) plus Gaussian noise with a standard deviation of 15 degrees. The mean motion orientation of the stimulus was adjusted on each trial so that participants performed consistently around 71% in an adaptive 2-up-1-down staircase procedure. This ensured that detection performance (*d’*) of all participants was roughly equal enabling a higher sensitivity for assessing metacognitive performance^23,35^. A full description of the motion stimuli and the staircase procedure can be found in Allen et al.^3^.

### Behavioural analysis

To assess performance of our groups we compared confidence ratings, accuracy, signal strength (stimulus motion orientation) and reaction times using independent-sample t-tests. To allow staircase stabilization, we discarded the first 30% of trials (three blocks total). Additionally, any missed trials were excluded from all analyses. Repeated-measures ANOVA confirmed that performance was stable (Fig. 1B) for the remaining seven blocks (F(6,222)=1.15, p=.337).

### Metacognition model

To assess the metacognitive ability of our participants we examined metacognitive efficiency *(M-ratio)*, which is a measure of how well one can consciously monitor their performance by controlling for the participant’s perceptual performance and task-independent biases in the confidence rating (for a full discussion see Fleming and Lau^23^). The *M-ratio* is calculated as the ratio between the metacognitive sensitivity *meta-d’* and the perceptual sensitivity *d’* ^35^. We focus on an *M-ratio* metric as it controls for potential perceptual differences between the groups^23^, and is important as OCD patients have previously been found to have worse perceptual decision making performance^24^.

To calculate an *M-ratio*, we used a hierarchical metacognition toolbox (hierarchical meta-d’ model HMM^36^, https://github.com/smfleming/HMM) as hierarchical methods are known to have a regularising effect and thus give more robust results. A detailed description of the model is found in Fleming^36^. In brief, it fits a signal-detection theoretic metacognition model to each group in a hierarchical fashion with group estimates of metacognitive efficiency (*m-ratio=meta-d’/d’*) governing individual participants’ behaviour. In particular, the model estimates a group-level metacognitive efficiency which governs individual subjects’ efficiencies. The optimisation of the metacognitive efficiency rather than the *meta-d’* was chosen because the latter is influenced by *d’*, which could bias metacognition results if *d’* were different^36^. By assessing metacognitive efficiency, this potential bias is taken care of^35^. We used the standard priors for the model, as derived from the developer’s previous studies^36^. Group parameters are estimated using Markov-Chain Monte-Carlo methods (MCMC, here: 3 chains of 10’000 samples each, burn-in of 1000 samples) implemented in JAGS (http://mcmc-jags.sourceforge.net). Model convergence was ensured by inspecting MCMC samples as well as checking that the *Ȓ* convergence measures for all parameters were <1.1 ^36^. We estimated each group separately and then compared the posterior group distributions in metacognitive efficiency. We followed this approach because our task was relatively short and individual parameter fits could be unstable or difficult to identify. To assess significance we computed the difference of the metacognitive efficiency group posteriors and compared the overlap with 0 of the resulting distribution (similar to an ordinary statistical test, it assesses the probability of the difference between the groups to be 0), as well as the 95% high density intervals of the difference distribution (analogous to confidence intervals).

### Perceptual decision making model

To explore differences in perceptual decision making between the groups and to test whether high compulsives showed similar impairments as patients with OCD^24^, we used an hierarchical version of a drift diffusion model^37^. The hierarchical drift diffusion model (HDDM^38^) estimates group model parameters using MCMC, similar to the metacognition model described above and has been used previously in investigating perceptual decision making in patients with OCD^24,39^.

Because we controlled for performance by adjusting signal strength (stimulus motion orientation), our analysis differed slightly from standard drift diffusion analyses. In our task, difficulty was directly related to the stimulus motion orientation, which changed on every trial. It is well known that stimulus strength directly influences the accumulation of evidence^40^, as it is depicted by the drift rate *v*. We thus used the motion direction on each trial to predict *v*.

To assess group differences, we entered both groups into the same hierarchical model, but used the group membership to predict differences in the model parameters. This deviates from the metacognitive analysis, in which we estimated both groups separately. We applied this approach, because the HDDM^38^ offers the possibility to explicitly model a group factor, a feature which is not yet implemented in the metacognition toolbox^36^. Such an approach can help to further increase the robustness of the parameter estimates^38^. We assessed four different models where (i) group influenced drift rate directly, (ii) group and orientation effect on drift rate interacted (i.e. group predicts how strong orientation affects drift rate), (iii) group has a separate effect on decision threshold *a*; and (iv) group interacted with orientation effect on drift rate as well as a separate group effect on threshold. Models were compared using deviance information criterion (DIC)^38^, and posterior group parameters of the best-fitting model were further assessed.

## Results

### Behavioural performance

To attain a stable proportion of correct and incorrect responses for all participants we adapted the difficulty of the dot motion paradigm (Fig. 1A) by adjusting the motion orientation of the stimuli using a staircase procedure. The groups thus did not differ in response accuracy (Fig. 1C; low compulsives: 73.54±2.54; high compulsives: 73.31±2.64; t(36)=0.28, p=0.780). Additionally, they did not differ in response latencies (Fig. 1D; low compulsives: 0.61s±0.12; high compulsives: 0.56s±0.11; t(36)=1.55, p=0.131). However, the stimulus motion orientation (median signal across trials), was significantly greater in high compared to low compulsive participants (Fig. 1E; low compulsives: 3.05 degrees±0.83; high compulsives: 4.25±2.05; t(36)=-2.42, p=0.021). High compulsive participants thus required a stronger motion orientation signal to perform at the same error rate as the low compulsives.

Comparing mean confidence rating we found no significant difference (Fig. 1F; low compulsives: 71.17±22.51; high compulsives: 68.56±19.64; t(36)=0.38, p=0.706). This means that high compulsive participants were neither more, or less, biased in reporting subjective confidence. Mean confidence, however, gives little insight into how well participants can consciously monitor their performance. To examine metacognitive differences between the groups, we thus used a hierarchical meta-d’ model.

**Figure 1.**
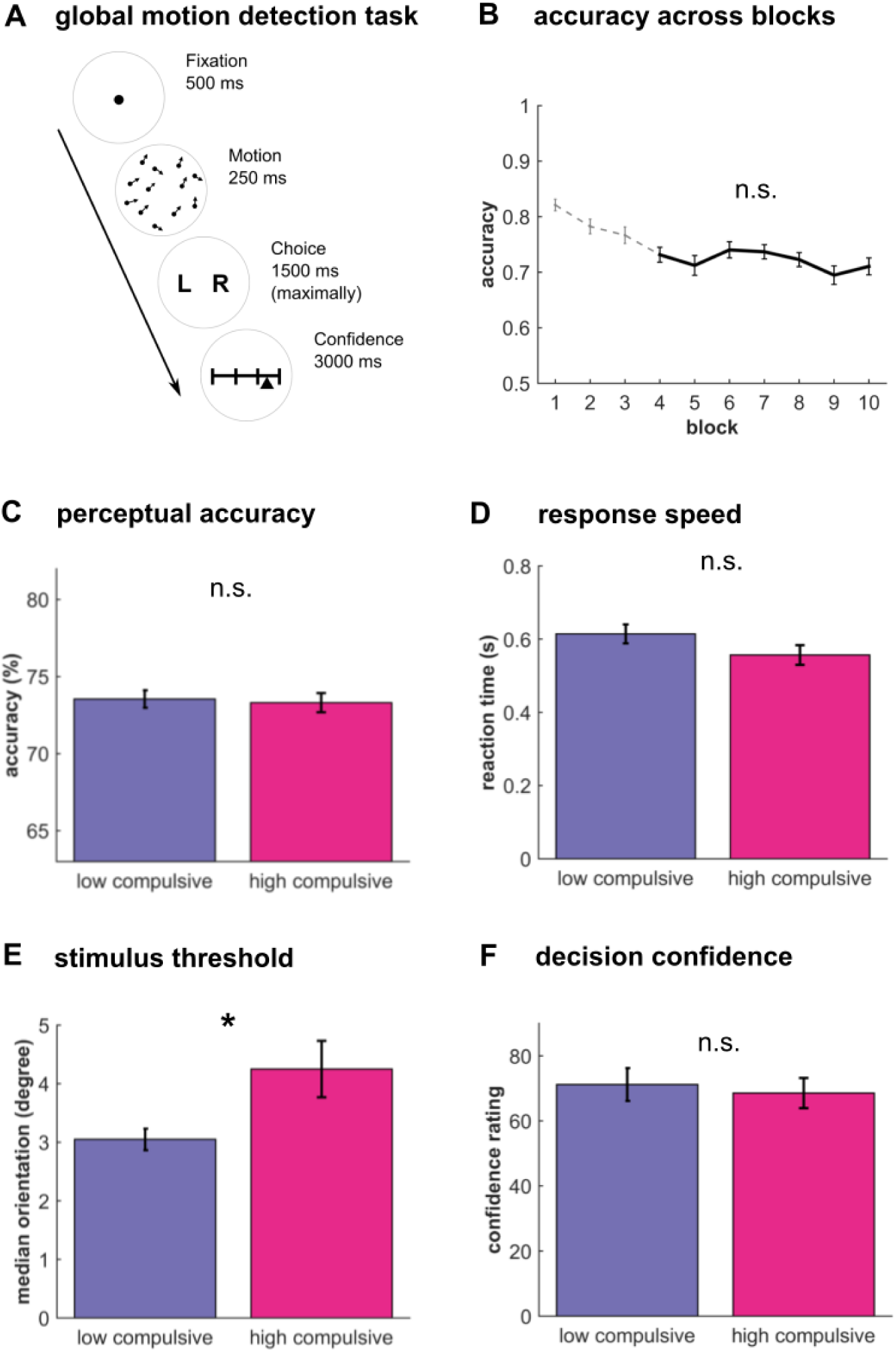
Metacognition task performance. High and low compulsive participants performed a metacognition task (A). Participants saw a cloud of dots moving with a defined mean motion orientation plus added random movement noise. After participants’ categorical judgement of the main direction of stimuli they then had to rate their confidence using a visual slider. (B) A staircase procedure ensured that performance was stable at 71% accuracy (the first three block were omitted (dotted line), because stability was not yet reached). This staircase ensured that both groups performed at the same level (C) and did not differ in their mean reaction times (D). Mean confidence ratings were similar between groups (F), but the sensory signal was significantly stronger in high compulsives (E), indicating a poorer perceptual decision making performance in high compulsive participants. Bar plots: mean±s.e.m; * p<.05; n.s. p>.05.

### Metacognitive impairments in high compulsive participants

For each group, we ran a separate hierarchical metacognition model^36^ and then compared the posterior group estimates for their metacognitive efficiency (*M-ratio*). This provides a measure of how well one can consciously monitor one’s own performance^23^. This signal detection theoretic measure equals 1 if an agent has full access to their perceptual performance, whereas values below 1 mean that confidence reports are suboptimal and cannot access full perceptual information. Computational modelling revealed that low compulsive participants have a mean metacognitive efficiency (*M-ratio*) of .814 (Fig. 2A, left panel), whereas high compulsive participants have a ratio of .512 (Fig. 2A, right panel). A comparison of the group posteriors revealed that the metacognitive efficiency was significantly lower in high compulsive participants (Fig. 2B; p(difference ≤ 0)=.017; equivalent to a one-sided significance test; 95% confidence intervals=.031-1.000). Importantly, this was not due to an impaired perceptual performance, as there was no significant group difference in their *d’* (Fig. 2C; t(36)=1.46, p=.153). These findings suggest that high compulsive participants have worse conscious access to their performance over and above any perceptual decision making impairments or response biases.

**Figure 2.**
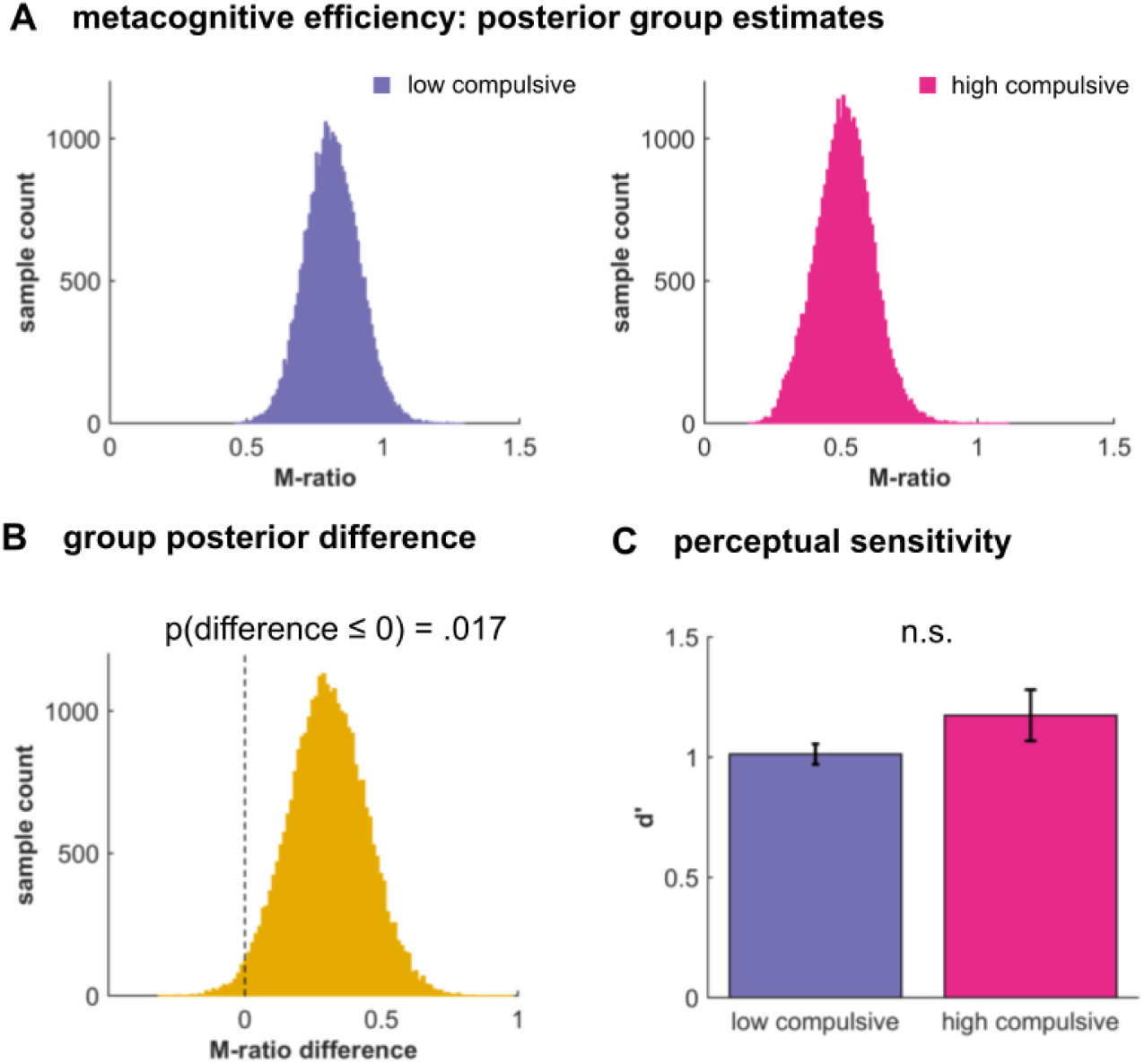
Metacognitive impairments in high compulsives. (A) Group posterior of metacognitive efficiency (M-ratio) for high and low compulsive participants revealed that high compulsive participants are significantly worse in their metacognitive abilities (B). This is not due to perceptual differences, because we controlled for performance, also indicated by the absence of a difference in the perceptual performance (d’, C). Bar plots: mean ± s.e.m; n.s. p>.10.

### Lower drift rate in high compulsives impairs perceptual decision making

A previous patient study found that OCD was associated with impaired perceptual decision making, especially with lower drift rates^24^. To explore the computational mechanisms causing the observed perceptual decision making impairments in high compulsive participants and to replicate and extend the previous findings, we applied a hierarchical drift diffusion model^38^. Model comparison (Table S1) revealed that the drift rate was modulated by task difficulty, as reflected in stimulus motion orientation. A model with a group factor (low, high compulsives) that modulates drift rate and its interaction with stimulus orientation, but not decision threshold, performed best.

To understand more precisely how the groups differ in their perceptual decision making, we evaluated the posterior model parameters of the best-fitting model. A highly significant influence of orientation on drift rate (Fig. 3A; p(*v_orientation_* ≤ 0)<.001) confirmed that stimulus difficulty directly influences evidence accumulation. The group factor had a highly significant impact on the relationship of stimulus orientation to drift rate (Fig. 3B; p(*v_orientation*group_* ≥ 0)<.001), meaning that high compulsive participants benefited less from the stimulus strength. The absence of a main effect of group on the drift rate suggests that there are no additional group-factors impacting the drift rate (Fig. 3B; p(v_group_ ≤ 0)=.091).

**Figure 3.**
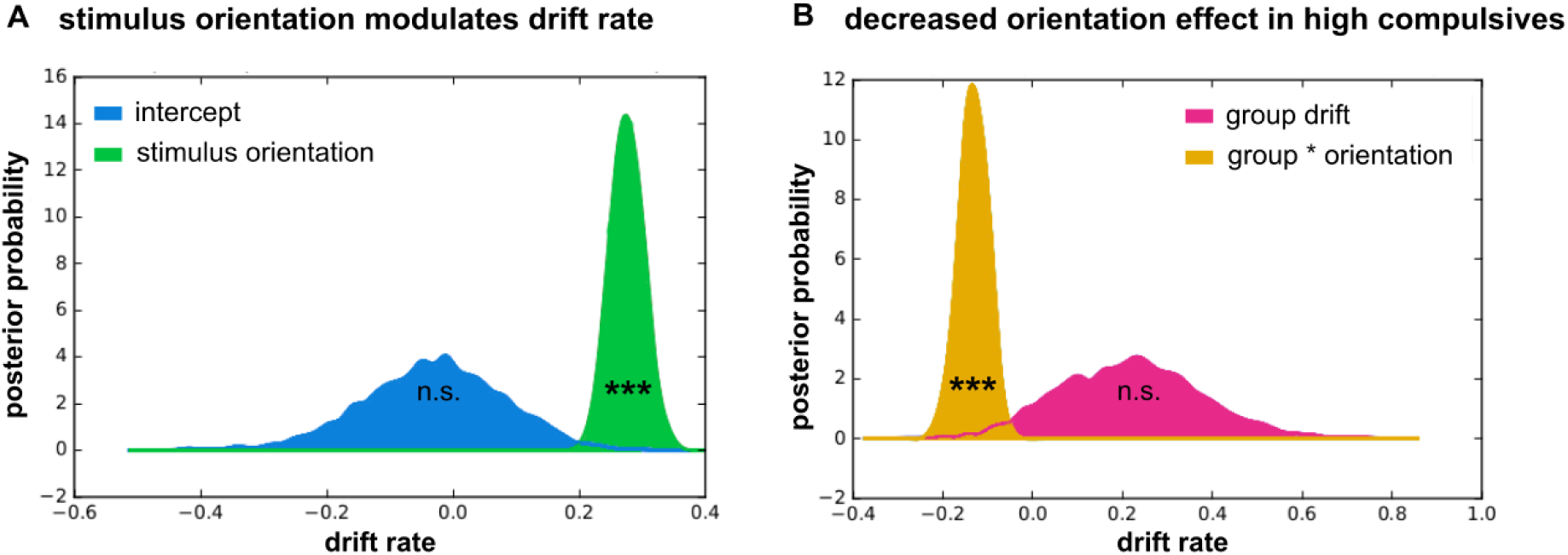
Stimulus processing is altered in high compulsive participants. (A) Signal strength (stimulus motion orientation) significantly increases drift rate across both groups (green). This effect entirely accounts for drift rate, as the orientation-independent drift rate (‘intercept’, blue) is not significantly different from 0. (B) The groups differ in in how much the stimulus motion orientation affects the drift rate: high compulsive participants benefit significantly less from an increasing stimulus orientation (orange). There is no additional effect of group on the drift rate (pink). *** p<.001; n.s. p>.05.

## Discussion

A longstanding tradition associates compulsivity with impairments in metacognition, but up to now metacognitive abilities have not been formally examined using computational models. In this study, we provide the first evidence that compulsivity is linked to an impairment in metacognitive abilities, independent of an additional impairment in perceptual decision making. This suggests that people with high compulsive traits are worse at introspectively monitoring their performance, and suffer from a degraded impact of sensory evidence on confidence.

Metacognition has traditionally been characterised as “thinking about thinking” or insight, a form of conscious monitoring of one’s own action^1^. Humans differ considerably in their metacognitive abilities^3,4^. Metacognitive investigations in OCD have mainly focused on biases in stimulus-outcome beliefs, for example that which an intrusive thought is likely to instantiate^9^. Later accounts have focused on memory-related confidence judgements, although with mixed results and heterogeneous approaches^10–15^, suggesting that OCD patients are not impaired in their memory, but in their confidence about their memory. Here, we expand on this research by showing that compulsive participants’ impairments are not restricted to biased beliefs or lowered confidence. Instead, we show that for high compulsive participants metacognitive judgements are less efficient, i.e. they are generally worse at accessing their own performance, a finding that holds when controlling for general response biases or perceptual decision making difficulty. This is of importance because it shows that compulsivity is related to impairments in metacognition, which sheds new light on the previous findings and theories. An impaired conscious access to one’s own performance can directly deteriorate the attitude towards intrusive thoughts and memories, as a poor monitoring system might induce a general distrust into one’s perceptions and recollections, and thus fosters distrust in memory recollection and an engagement in compulsive safety behaviours.

As reported in OCD patients^24,41^, we found that high compulsive participants have perceptual decision making impairments in the visual domain. This was expressed in our task by an increased stimulus motion orientation (i.e. signal strength), and our computational modelling related this impairment to a lower drift rate, in accord with this previous study^24^. It is interesting to speculate how this perceptual decision making difficulty might be related to the metacognitive impairments. At its most simple, these impairments could be completely independent, so that compulsivity is contributed to by a lower metacognitive efficiency as well as a lower perceptual decision making sensitivity. Alternatively, perceptual decision making impairments could indirectly affect metacognition in a bottom-up manner by also influencing a post-decision evidence accumulation process^42–44^. However, it is unclear how the increased signal strength for high compulsives would influence a post-decision accumulation. Lastly, a perceptual decision making difficulty could be a top-down consequence of impaired metacognition, where impaired metacognition alters the amount of evidence a participant needs to make a decision. This in turn could impact their behaviour in perceptual decision making tasks, such that they only decide once they have consciously perceived enough information, leading to an increased need for greater signal strength.

We focused on ‘healthy’ participants, selected form a large population-based sample, who scored high or low on a compulsivity scale. This had the advantage of controlling for psychiatric dimensions that are often comorbid with compulsivity, such as depression and anxiety. This is important given that metacognitive impairments are suggested to be symptomatic for many psychiatric disorders^45^. Thus, our experimental strategy allows us to be confident that observed differences are solely driven by compulsivity, but not by other psychiatric traits. Additionally, our replication of a perceptual decision making impairment similar to the one found in patients with OCD^24^ speaks to a conceptualisation of compulsivity in terms of as a spectrum, rather than as a categorical entity^25,26^. However, future studies of patients with OCD will be necessary to ascertain whether similar processes are impaired in participants with clinically relevant compulsivity.

In summary, we show that a compulsivity spectrum identified in the general population is linked to impairments in metacognitive efficiency. This impairment is expressed over and above an effect due to perceptual decision making difficulty. Our findings provide the first computational evidence that metacognition is impaired in compulsivity and thus clarify the relationship between compulsivity, perceptual performance and conscious insight.

## Author contributions

TUH, MA, GR & RJD designed the study. TUH & NSPN consortium collected data. TUH & MA analysed the data. RJD supervised data collection. TUH, MA, GR & RJD wrote the paper. NPSN consortium provided database for recruitment. All authors approved the final version of the manuscript.

## Acknowledgements

We thank Liz Harding, Danae Kokorikou, Hina Dadabhoy, Alexandra Hopkins, and Kalia Cleridou for helping with the recruiting. We thank Gita Prabhu for her support with the ethics application, recruiting and data organisation of this study. We also thank the NSPN Consortium for providing access to the UCHANGE database, and Michael Moutoussis for his help with the database handling. We thank Steve Fleming for his support for the metacognition modelling, and Rani Moran on his comments on an earlier draft. A Wellcome Trust Cambridge-UCL Mental Health and Neurosciences Network grant (095844/Z/11/Z) supported RJD and TUH. RJD holds a Wellcome Trust Senior Investigator Award (098362/Z/12/Z). GR and MA were supported by a Wellcome Trust SRF grant (100227). The UCL-Max Planck Centre is a joint initiative supported by UCL and the Max Planck Society. The Wellcome Trust Centre for Neuroimaging is supported by core funding from the Wellcome Trust (091593/Z/10/Z).

